# Compact bifurcation analysis of the Epileptor

**DOI:** 10.1101/2023.10.26.564167

**Authors:** Maria Luisa Saggio, Viktor K. Jirsa

**Affiliations:** Institut de Neurosciences des Systémes - Inserm UMR1106, Aix-Marseille Université, Marseille, France

## Abstract

The Epileptor is a phenomenological model able to reproduce the activity of the most common class, in terms of dynamics, of epileptic seizures, characterized by having square-wave bursting properties. It also encodes an additional mechanism to account for interictal spikes and spike and wave discharges. This model is being used in large-scale brain modeling of epileptic patients with the goal of improving surgical outcomes. Here we use insights from a more generic model for square-wave bursting, based on the Unfolding Theory approach, to guide the bifurcation analysis of the Epileptor. This allows to understand how the Epileptor’s parameters can be modified to produce activities for other seizures classes (i.e. other onset/offset bifurcation pairs) as observed in patients and to unveil how the interaction with the additional mechanism for spike and wave discharges alters the bifurcation structure of the main burster.

## 1 Introduction

Epilepsy is the most common among the chronic and severe neurological diseases, affecting 65 million people worldwide [Moshé et al., 2015] and the complexity of this group of disease unfolds along different axes. The complexity comprises its multifactorial causes, since ‘almost any condition affecting the cerebral grey matter can result in epilepsy’ [Shorvon et al., 2009], and the presence of interacting processes spanning several scales in time (from mechanisms producing very fast 2kHz oscillations to ultra-slow processes spanning the entire lifespan of an individual) and space (from within neuron processes to brain regions and whole brain network mechanisms). A variety of pathological processes leads to seizures that are remarkably similar in their electrographic signature [Jirsa et al., 2014] because, physiologically, they all lead to the instability of gray matter as stated already in 1874 by Hughlings Jackson [Taylor, 1930].

For these reasons phenomenological models are particular useful in the study of epileptic seizures [Saggio and Jirsa, 2022], because they allow to investigate the dynamical mechanisms related to this instability without committing to a specific choice in terms of biological implementation. Understanding the dynamical mechanisms underlying seizure activity could potentially help devise strategies to exit from the ictal state and abort a seizure, or to prevent the system to enter the ictal state altogether [Brogin et al., 2020; Izhikevich, 2007; Szuromi et al., 2023]. In addition, due to their lower computational burden, phenomenological models for seizures activity are prime candidates to be used in whole-brain simulations.

Insights from this class of models can also be used to better understand the dynamic repertoire of other models with a wider range of parameters, as typically occurs for bio-physically inspired ones, establishing a link between dynamical and physiological mechanisms (see for example Depannemaecker et al. [2022]). Simpler phenomenological models, though, can also guide the investigation of other more complex phenomenological ones.

In this article, we use a generic model for bursting activity, for which we have a deep understanding of the dynamical repertoire, to guide the analysis of the more complex, but still phenomenological, Epileptor model.

The Epileptor model has been proposed by Jirsa et al. [2014] to reproduce the most predominant class of seizures as observed in *in vitro* preparations, zebrafish, mice and human epileptic patients. The class is defined as the onset/offset bifurcations pair that delimitates a seizure [Jirsa et al., 2014], extending to epilepsy a taxonomy developed by Izhikevich [2000] and mainly applied to neuronal bursting. The most common class for seizures identified in data had Saddle-Node (SN) bifurcation for the onset of the fast oscillations and a Saddle-Homoclinic (SH) bifurcation for their offset [Jirsa et al., 2014]. The Epileptor phenomenologically incorporates this fast-slow dynamics, together with other forms of epileptiform activity, such as preictal spikes and spike-and-wave complexes. It has been noted that a minimal model for this type of bursting (fast-slow SN/SH bursting, also known as square-wave bursting) can be obtained by using a layer of the unfolding of the degenerate Takens-Bogdanov (DTB) singularity to create a ‘map’ of possible behaviors and by adding a slow dynamics to promote movement on this map [Golubitsky et al., 2001] and that this layer could host other types of bursters [Osinga et al., 2012]. Building on this, we have developed a minimal model with a rich repertoire of classes from the taxonomy [Saggio et al., 2017]. We will refer to it as the DTB bursting model.

In the present work we use insights from fast-slow bursters and the DTB bursting model to investigate the Epileptor model. In particular, we identify the fast parameters that contain the parametrization, in terms of slow variables, of the ‘path’ through which the fast subsystem is pushed. We explore the bifurcation diagram of these fast parameters (‘the map’) to identify all the relevant bifurcation curves we expect to find based on the knowledge of the unfolding of the deg. DTB singularity [Dumortier et al., 1991]. We show how the Epileptor moves on the map during a seizure and how alternative paths with different onsets and offsets (as observed in patients [Jirsa et al., 2014; Saggio et al., 2020]) can be placed in this single map, some of them not yet observed in the Epileptor. We highlight how the input from the additional system for preictal spikes and spike-and-wave complexes alters the path on the map, and thus the sequence of bifurcation curves encountered by the system. Finally, we show the effect that the main Epileptor’s parameter, the epileptogenicity *x*_0_, that is the brain region proneness to seizures, has on the path.

The Epileptor is currently the model of choice in the Virtual Epileptic Patient (VEP) framework, a patient-specific large-scale model of brain activity with the potential of helping presurgical evaluation of drug resistant patients [Jirsa et al., 2017; Proix et al., 2017]. For some of these patients, an alternative to medication is the surgical resection of the brain regions involved in the generation of seizures, the epileptogenic zone (EZ), under the constrain of limiting post-surgical neurological impairments [Bartolomei and Wendling, 2009]. However, outcomes of this type of surgery are very variable and depend on the patient condition and epilepsy, with rates of success ranging between 34% and 74% [Jobst and Cascino, 2015]. The inference of the EZ is highly non trivial and the VEP approach wants to arm clinicians with an additional tool to use in their evaluation. This tool leverages on the possibility, through whole brain simulations, to reveal otherwise hidden complex dynamical effects. It can be used to test specific clinical hypotheses (e.g. ‘will seizures stop by removing a specific set of brain regions?’) by simulating functional data or to find unbiased optimal surgery strategies through parameters fitting techniques.

Patient-specificity is key and, under the requirement that state of the art methods are used in diffusion imaging, it has been shown that the patient-specific connectome gives the best outcome for the VEP [Proix et al., 2017]. However, at the brain region level, one single type of bursting class, the SN/SH of the Epileptor model, is being used for all patients. This despite the fact that patients data show heterogeneity in the bursting class [Jirsa et al., 2014; Saggio et al., 2020] and that different classes may behave differently in terms of synchronization and propagation properties [Izhikevich, 2007] so that the class choice can potentially alter the VEP outcome.

For these reasons, understanding the full potential of the Epileptor model in terms of bursting dynamics and how to set the parameters in order to change class, is an important step to improve the patient-specificity, and potentially the predictive power, of VEP models.

Beyond surgery, a promising avenue to terminate seizures is that of brain stimulation. In silico experiments through the VEP framework could also include the design of patient specific stimulation protocols to prevent or abort seizures [Jirsa et al., 2017]. Here, as well, given the different ways in which bursters react to stimulation [Izhikevich, 2007], patient-specificity in terms of bursting class could prove to be crucial [Szuromi et al., 2023].

## 2 Models

### 2.1 The DTB bursting model

The DTB bursting model [Saggio et al., 2017] uses the unfolding of the DTB singularity as fast subsystem, this gives a map of the possible behaviors when the unfolding parameters (*μ*_1_, *μ*_2_, *ν*) are modified. The portion of this map that is relevant for SN/SH bursting can be obtained by taking a layer of the unfolding for fixed positive *μ*_2_ or for fixed positive *ν*. If the fixed parameter is chosen small enough, the maps obtained will be topologically equivalent [Dumortier et al., 1991]. Here we consider a layer for *μ*_2_ = 0.07, as shown in Fig 1A. There are regions with only one stable attractor, either a fixed point or a limit cycle (white) and regions of bistability between two fixed points (grey) or a fixed point and a limit cycle (yellow). These regions are separated by bifurcation curves: SN, SH, supercritical Hopf (SupH) and Saddle-Node-on-Invariant-Circle (SNIC).

**Figure 1:**
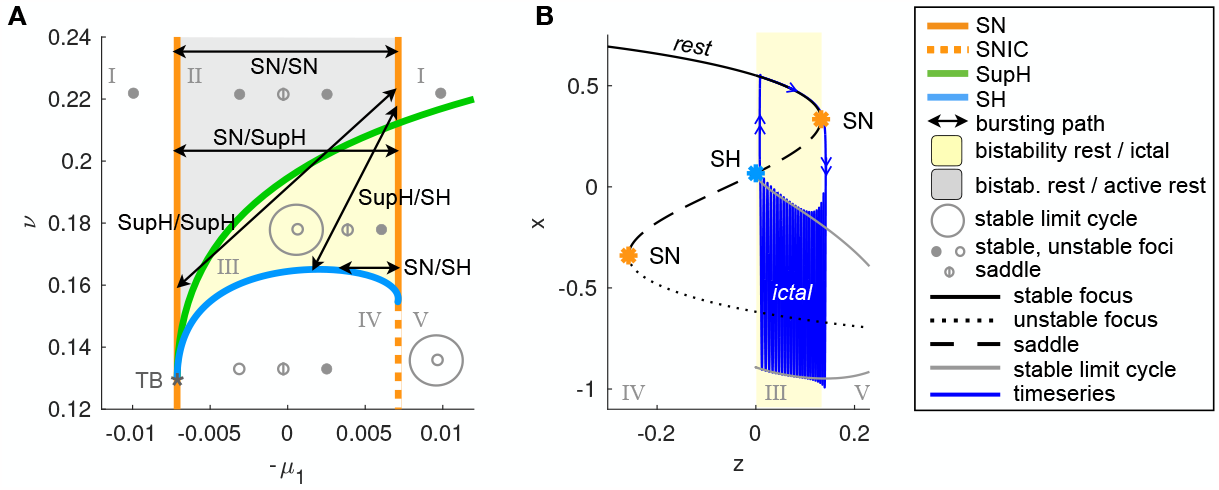
Hysteresis-loop bursting in the DTB bursting model. **A**. This is one portion of the unfolding in which SN/SH bursting can be placed, together with other classes [Saggio et al., 2017]. Saddle-Node (SN) and supercritical Hopf (SupH) curves meet at the Takens-Bogdanov point TB. Bifurcation curves partition the map in five regions with different state space configurations (Roman Numerals). When this map is used for the fast subsystem of a fast-slow bursters with an hysteresis-loop mechanism for the slow variable, possible classes in the map are: SN/SH, SN/SupH, SupH/SH and SupH/SupH plus SN/SN where the system alternates between the two stable fixed points [Saggio et al., 2017]. When more than one fixed point exist, the resting (or inter ictal) state is the one on the right, the other one we call ‘active rest’. The resting state corresponds to the upper branch of fixed points in panel B. **B**. Typical bifurcation diagram for the SN/SH class. When the system is at rest, *z* increases until the fixed point destabilizes through the SN bifurcation and the system jumps into the stable limit cycle. Now that the system is far from rest, *z* decreases until the limit cycle destabilizes through a SH bifurcation and the system jumps back to rest. If the destabilization of the fixed point/limit cycle is obtained through a different bifurcation, we will have a different onset/offset class and a different appearance of the burster’s timeseries.

If the two parameters (*μ*_1_, *ν*) slowly change as a function of a variable *z* we can have movement on the map. We parametrize (*μ*_1_(*z*), *ν*(*z*)) so that the allowed movements are straight lines:

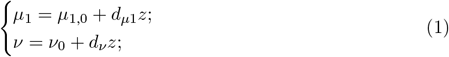

with (*μ*_1,0_, *ν*_0_) being the initial point of the path and (*d*_*μ*1_, *d*_*ν*_) the direction vector of the path. We then impose a simple dynamics for *z* such that: when the system is at or close to rest *z* increases and the fast subsystem moves rightward on the map; when the system is far from rest *z* decreases and the system moves leftward.

The model equations read:

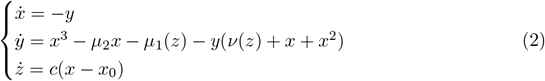

where *c* ≪ 1 gives the time separation among the two subsystem and *x*_0_ sets the threshold for inverting the behavior of *z*. In this types of models *x*_0_ is called ‘excitability’, because it expresses how prone the system is to move towards the destabilization of the resting state. In Eq (2) the dynamics of the slow variable has been simplified as compared to the original model, in a way that is only possible in some regions of the unfolding. See the Methods Section for more details.

If we initialize our fast subsystem inside the bistability region, when at rest it moves towards a SN curve that destabilizes the fixed point and forces the system to jump to the limit cycle (or to the other fixed point if in the grey region) now the fast subsystem starts moving leftwards until it reaches a bifurcation that destabilizes the new attractor so that the fast subsystem jumps back to rest. This loop is what constitutes fast-slow hysteresis-loop bursting. Depending on where the path is placed on the map, the system will encounter specific sequences of bifurcation curves, so that different types of bursting are possible (Fig 1A).

Since this model is generic for SN/SH bursting, we can expect that a topologically equivalent map exists in the Epileptor model for some values of the parameters and that it is possible to adjust these parameters to have other bursting classes.

### 2.2 The Epileptor

For the Epileptor model, we use equations as in Jirsa et al. [2014], with the addition of the parameter *m* as in El Houssaini et al. [2015]:

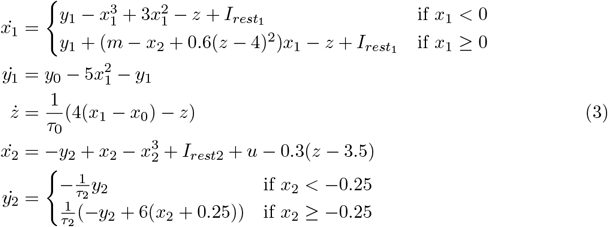

where

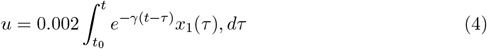

with *x*_0_ = −1.6, *y*_0_ = 1, *τ*_0_ = 2857, *τ*_2_ = 10, *τ*_1_ = 1, 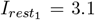, *I*_*rest*2_ = 0.45 and *γ* = 0.01.

The Epileptor is composed of three subsystems acting on different time scales: a fast, an intermediate and a slow subsystem.

A fast subsystem (*x*_1_, *y*_1_), acting on the scale described by the time constant *τ*_1_, is based on a modified Hindmarsh and Rose model and is responsible for the fast oscillatory activity observed during a seizure. This system can display bistability between a stable fixed point (rest or interictal condition) and a stable limit cycle (fast oscillatory activity or ictal state), similarly to the yellow region in Fig 1A. As in the DTB bursting model, the slow variable *z* allows for hysteresis-loop bursting in this region (Fig 2, blue). This mechanism is the core of the Epileptor model.

**Figure 2:**
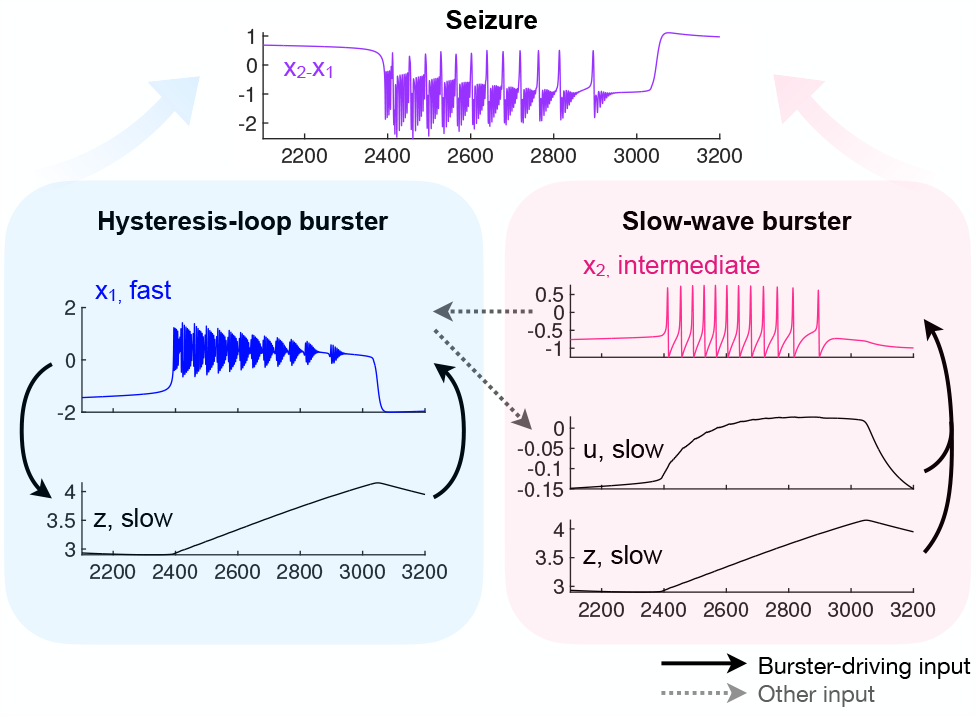
The Epileptor’s main mechanisms. The Epileptor field potential (purple), is a combination of the activity of one fast variable (blue) and an intermediate one (magenta). The fast and slow variables dynamics and feedback among them constitute the core of the model (blue), while the intermediate variables (magenta) modulate the fast activity. All the simulations in this work are performed without noise.

In addition, subsystem (*x*_2_, *y*_2_) acts on an intermediate time scale, given by *τ*_2_, and is an excitable system with a SNIC bifurcation. It is responsible for the generation of preictal spikes, which reflect the increased excitability of this subsystem close to seizure onset, and for spike and wave complexes during the seizure. It displays ‘slow-wave’ SNIC/SNIC bursting (Fig 2, magenta) in which the role of slow variables is played by *u*, a low-pass filtered input from *x*_1_ (see Eq (4)), combined with *z*. In this type of bursting there is no need for bistability nor feedback between the faster (i.e. the intermediate system) and slower variables [Izhikevich, 2000]): independently from what the intermediate variable is doing, the slow variable pushes it back and forth across a SNIC bifurcation. The activity of the intermediate subsystem modulates that of the fast one.

The Epileptor field potential (Fig 2, purple) is given by a combination of fast and intermediate variables: *x*_2_ − *x*_1_.

## 3 Results

Given the presence of time scale separation, with *τ*_1_ ≪ *τ*_2_ ≪ *τ*_0_, the three subsystems can be analyzed in isolation [Rinzel, 1985].

We here focus on the core hysteresis-loop mechanism. To do so we will study the relevant parameter space of the fast subsystem (this subsystem is the one containing the bifurcations for the onset and offset of seizures) to create a ‘map’ of its dynamical repertoire, guided by what we know about the map of the DTB bursting model. We will then follow the path of the full Epileptor model in this map.

### 3.1 Fast subsystem - The map

In this section we define, for better readability (*x*_1_, *y*_1_) = (*x, y*). We are interested in whether different types of behaviors can be produced when the fast parameters are allowed to vary. We thus rewrite the fast subsystem in Eq (3), to highlight the presence of the parameters of the model, as:

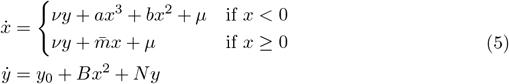

Here we consider *ν, a, b, μ*, 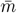, *y*_0_, *B* and *N* as parameters of the fast subsystem, and interpret

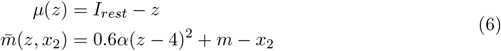

as a parametrization of the path that the fast subsystem follows in its parameter space as promoted by feedback from the slower variables *z* and *x*_2_.

The fast subsystem as in Eq. (3) can be obtained from Eq. (5) and (6) by replacing: *ν* = 1, *a* = −1, *b* = 3, *μ* = *I*_*rest*_ − *z, B* = −5, *N* = −1 and *α* = 1.

Since 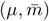 are the fast parameters that slowly change in the Epileptor’s dynamics, we expect that, when they are treated as bifurcation parameters, they will produce a map similar to that of the generic model. This entails, beside curves for the onset (SN) and offset (SH) bifurcations, the presence of a curve of H bifurcation that encounters the SH curve on a SN curve different than the onset one.

In all the figures and simulations we set parameters other than 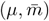 as in Eq (3), unless otherwise specified, with the exception of *x*_0_ = −2, which we have modified for visualization purposes. As explained later, this doesn’t alter the onset/offset pattern. However, in the bifurcation analysis, we will maintain all the parameters in order to gain some insights on their role in shaping the map.

#### 3.1.1 Bifurcation analysis

##### Fixed points

The fixed points of the system can be obtained by imposing:

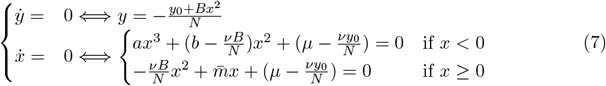

The solutions are such that, in the 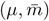 space, there is a central region with three fixed points, while outside this region there exist only one solution: the lower branch of the fixed points manifold for small values of *μ* and the upper branch for big values (Fig 3).

**Figure 3:**
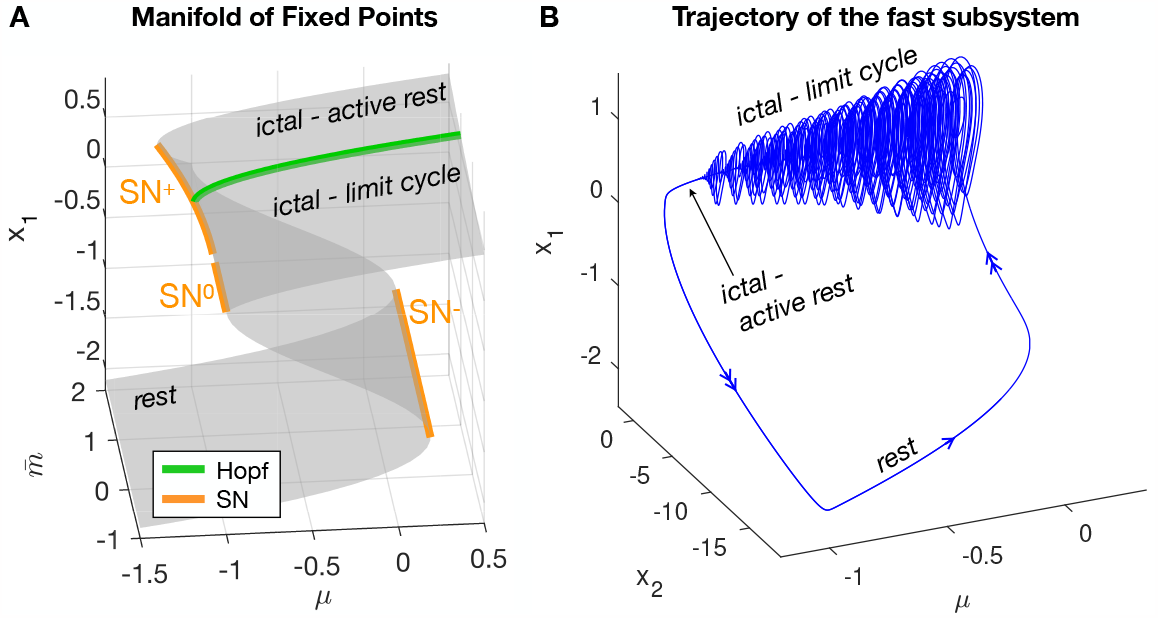
Epileptor’s fixed points and bursting trajectory. **A**. Fixed points manifold for *x*_1_. For *x*_1_ *<* 0, there are two solutions, while for *x*_1_ ≥ 0 we have one fixed point when 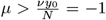 and two solutions for smaller values. The lower branch of fixed point is the rest or interictal state. The upper branch is stable for values of 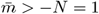 and unstable otherwise. *SN* ^+^ occurs for 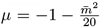, *SN*^−^ for 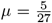, *SN* ^0^ for *μ* = −1 and the Hopf curve for 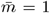. **B**. Trajectory of the Epileptor’s fast subsystem (*x*_1_, *x*_2_) plotted against the parameter *μ*. It can be observed the hysteresis-loop caused by the two *SN* curves.

##### Saddle-Node manifolds

The central region with three solutions is delimited by two curves of SN bifurcations, satisfying the additional condition that the determinant of the Jacobian of the system is zero:

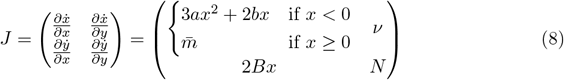

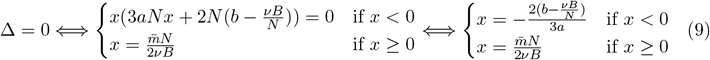

This gives a curve for negative values of *x, SN*^−^ and a curve for positive values of *x, SN* ^+^. When 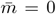 the positive portion of the curve ends because *x* = 0. It joins, however, with a curve we find for *x* = 0, where, for 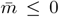 the positive and negative branches of fixed points merge. This is not a real SN bifurcation curve because it occurs where the system is piece-wise and the positive and negative limits of the Jacobian do not match. However, for the goal of bursting it will behave as a SN curve. With these caveats we call it *SN* ^0^.

By inserting the solutions in Eq (9) in Eq (7) we can find conditions on the parameters for the *SN*^−^ and *SN* ^+^ curves. Imposing *x* = 0 in Eq (7) we find that one for *SN* ^0^:

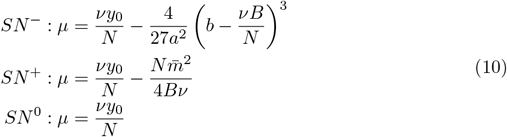

##### Andronov-Hopf manifold

Beyond being fixed points, candidate Hopf bifurcation points must satisfy the condition that the trace of the Jacobian should be equal to zero:

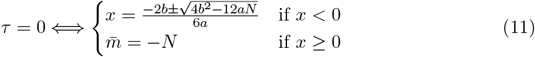

When inserting Eq (11) in Eq (7) we find no solutions for *x <* 0. For *x* ≥ 0, the condition 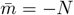 gives Δ *>* 0, and thus a Hopf curve [Dercole and Rinaldi, 2011], only on the upper branch of fixed points as shown in Fig 3A.

##### Codimension 2 Takens-Bogdanov manifold

We find a manifold of codimension 2 TB bifurcations where the Hopf and *SN* ^+^ merge, that is for 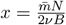.

##### Saddle-Homoclinic curve

Once identified the local bifurcations, we searched for the SH curve. Since this is a global bifurcation, we used numerical tools and we performed our analysis fixing all parameters as in Jirsa et al. [2014] except for those used for the map. It was problematic to locate it using continuation softwares (Matcont) possibly due to the system being piece-wise. We thus performed simulations for different values of the parameters 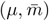, initializing the system close to the upper fixed point (either active rest or unstable fixed point) and computing the amplitude and frequency of the limit cycle with regards to the *x* variable (Fig 4 right panels). The trend of the frequency behavior is compatible with the presence of a SH bifurcation, which requires the frequency to scale to zero towards the bifurcation point. This curve stems from the TB point, as predicted from the theory, and reaches the other SN curve as in the unfolding of the DTB singularity. We have thus found a region of topological equivalence between the two models (Fig 4).

**Figure 4:**
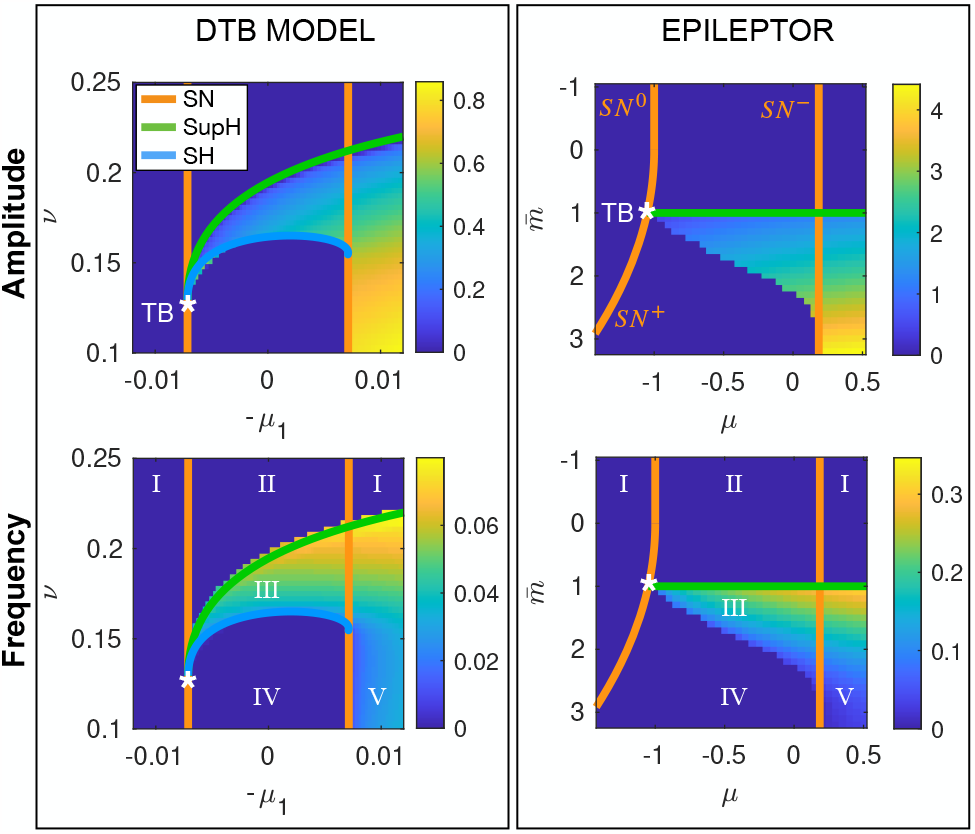
Topological equivalence between bifurcation diagrams of the fast sub-systems of the Epileptor and of the DBT bursting model. In the left panels we show the portion of bifurcation diagram of the DTB bursting model (that is the unfolding of the DTB singularity) in which SN/SH bursting occurs, and the behavior of amplitude and frequency of the limit cycle. In the right panels, the same for the Epileptor’s fast sub-system. In the latter case the presence of a SH bifurcation stemming from the TB point can be inferred by the behavior of the frequency (Hz) of the limit cycle identified through simulations, which scales down to zero. Roman Numerals refer to the configurations as in Fig 1.

##### Role of the other parameters on the topology of the map

We can appreciate the role of some parameters, other than 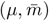, in shaping the map. For example, in Eq (10) we can see how they contribute to the SN curves. In particular, *y*_0_ pushes all the curves right or left on the map, while *b* and *a* only act on the negative branch. With regards to the position of the Hopf curve, it simply depends on *N*. We can’t make similar considerations for the SH curve, that we obtained numerically, except that the location of its starting point will depend on the location of the TB point where the H and the *SN* ^+^ curves meet.

Of course, these are considerations that only hold for small changes of the mentioned parameters. Understanding the intervals in which these parameters could be changed without altering the topology of the map would give information about its robustness. While useful, given that the physiological correlates of these parameters could fluctuate, this is beyond the scope of this work.

### 3.2 Slower variables - Paths on the map

Now that we have the bifurcation diagram for the slowly changing parameters of the fast subsystem, we can go back to the path followed by the fast subsystem on this map under the influence of the fast and intermediate variables. The path is parametrized as in Eq (6), which we rewrite here for convenience:

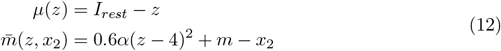

#### 3.2.1 Placing on the map classes from the literature

Movement parallel to the *μ* axes is promoted by *z* and is the core of the hysteresis-loop bursting, in which the slow variable, with feedback from *x*_1_, pushes the fast subsystem across the onset and offset bifurcations. By changing the parameter *m* we can move the path upward or downward, so that it will cross different pairs of bifurcation curves. In Fig 5, we plotted on the map the simulated paths for values of *m* from the literature [El Houssaini et al., 2015; Jirsa et al., 2014] and show how they produce bursting of different classes, namely SN/SupH, SN/SH and SN/SN (in the latter burster, the onset/offset do not refer to oscillations, but simply to the alternation between the fixed points in the upper or lower branches). Of note, for *m* = 0 as in the original Jirsa et al. [2014], input from *x*_2_ is such that the actual offset of the hysteresis-loop burster is a SupH rather than a SH (Fig 5A). The SH offset can be retrieved by setting *I*_*rest*2_ = 0 so that there is no bursting in the intermediate subsystem (not shown).

**Figure 5:**
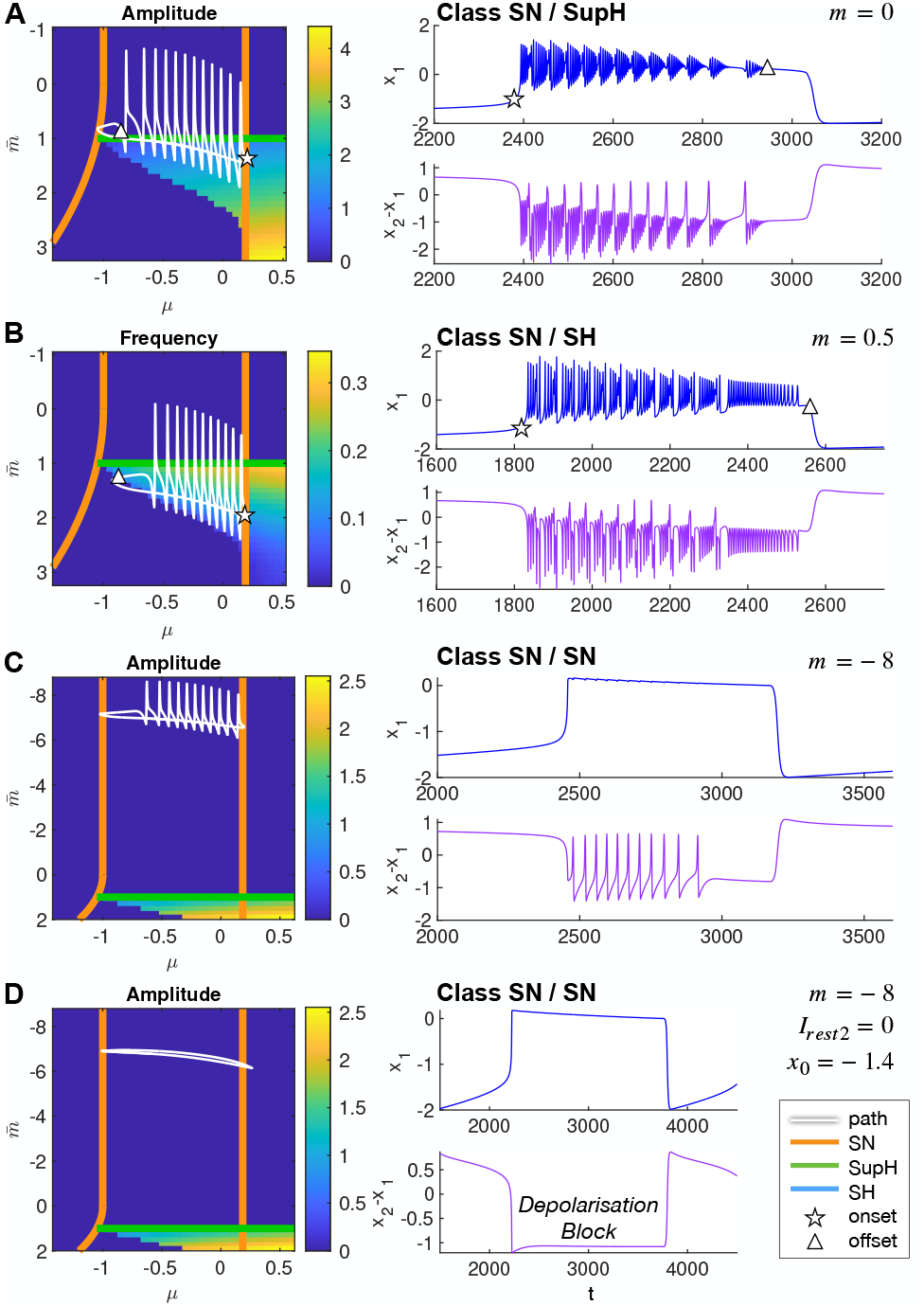
Paths on the map. Paths followed by the full Epileptor model in the map for four different conditions (left column), and the related timeseries (right column). All parameters are kept the same except for those specified. Both on maps and in timeseries, a triangle/star approximately marks the onset/offset of oscillations in the fast subsystem, when present. This types of Epileptor behaviors have been described in the literature.

#### 3.2.2 Intermediate subsystem modulatory effects

While the fast subsystem is in the ictal state (either oscillating or in the active rest), its low-pass filtered activity and input from *z* cause oscillations in the intermediate subsystem through a SNIC bifurcation. The intermediate fluctuations of *x*_2_, in turn, modulate the 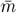 component of the path, and appear in Fig 5A-C as ‘spikes’. These spikes can cause the fast subsystem to cross the Hopf curve multiple times while moving towards seizure’s offset. This is visible in the amplitude changes in the timeseries, as the amplitude decrease when approaching this bifurcation (Fig 5A-B). For example, let’s consider the timeseries in Fig 5A. In the first part of the ictal state the fast subsystem briefly crosses the H curve but not long enough to settle down to the fixed point, while in the latter part of the timeseries the sequence of SupH bifurcations becomes more evident.

These ‘spikes’ in the path come from the activity of the intermediate subsystem, implying they are present even when *m* is chosen such as to bring the path fully above the bistability region as in Fig 5C. However, since in this region no fast oscillations are possible, their modulatory effect is negligible. Finally, by setting *I*_*rest*2_ = 0 so that the intermediate subsystem doesn’t reach the threshold for SNIC, as in Fig 5D, the spikes are no longer present in the path. We show it for *m* = −8. In this class, the active state of the Epileptor has been suggested to be linked to depolarization block [El Houssaini et al., 2015], a physiological state in which action potentials cannot be triggered despite the neuronal membrane being depolarized.

#### 3.2.3 Identifying new classes of hysteresis-loop bursting

The DTB bursting model allows for other two classes of hysteresis-loop bursting to be found in this map: SupH/SH and SupH/SupH (Fig 1A). By looking at Fig 5, we can observe that, in the Epileptor model, they can’t be obtained by a simple vertical translation of the path. However, by changing the global slope of the path setting *α* = −1 in Eq (6) and changing *m* to appropriate values, we could simulate these other two classes as shown in Fig 6.

**Figure 6:**
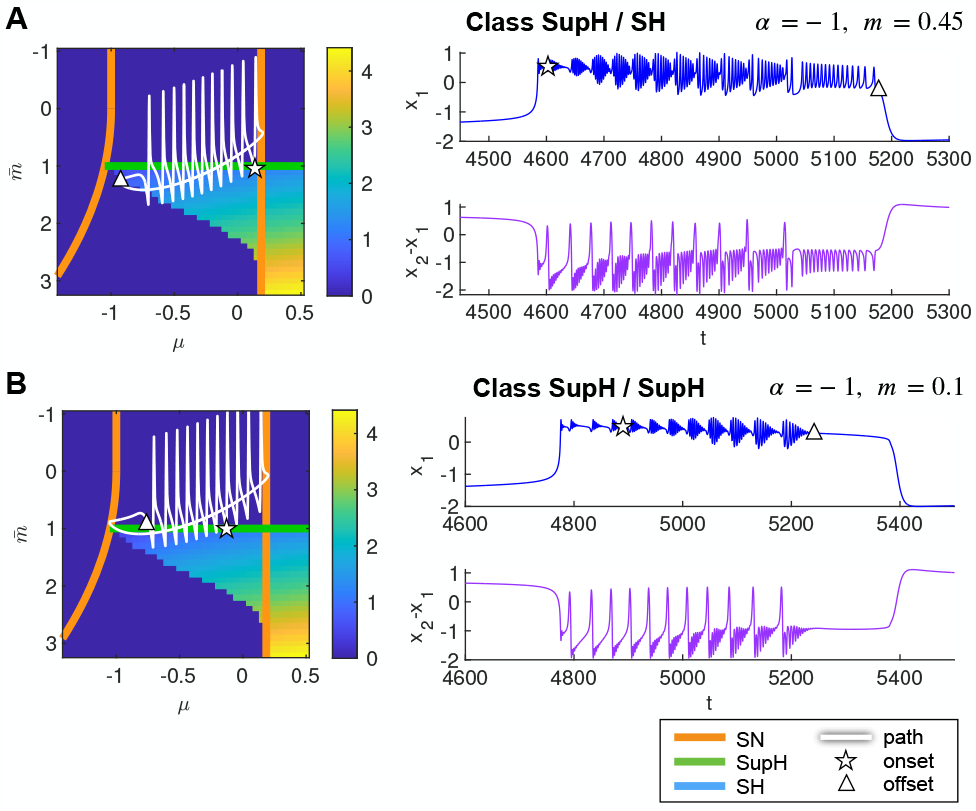
New classes of bursting identified in the Epileptor model. Path on the map (left) and timeseries (right) for the new classes SupH/SH (**A**) and SupH/SupH (**B**). Maps represent the amplitude of the limit cycle.

#### 3.2.4 Role of *x*_0_ on the path

With regards to the slow variable dynamics (Eq (3)), the relevant parameter is *x*_0_, that is linked to how close the z-nullcline is to the resting state (Fig 7A). As in the DTB bursting model, the closer the z-nullcline is to the resting state, the more slowly *z* evolves when at rest towards the ictal state and the faster it evolves when in the ictal state towards seizure offset (Fig 7B left, middle). When the z-nullcline crosses the branch of resting state, the intersection is a fixed point for the whole system (Fig 7B right) [El Houssaini et al., 2015]. In the VEP model, *x*_0_ is used to tune each brain region’s epileptogenicity, that is its proneness to start a seizure. When considering only the hysteresis-loop mechanism, this parameter doesn’t affect the path on the map, but rather the speed of movement along different parts of it. However, when the intermediate subsystem is in the oscillatory regime, different values of *x*_0_ allow for a different amount of intermediate spikes along the path (Fig 7C).

**Figure 7:**
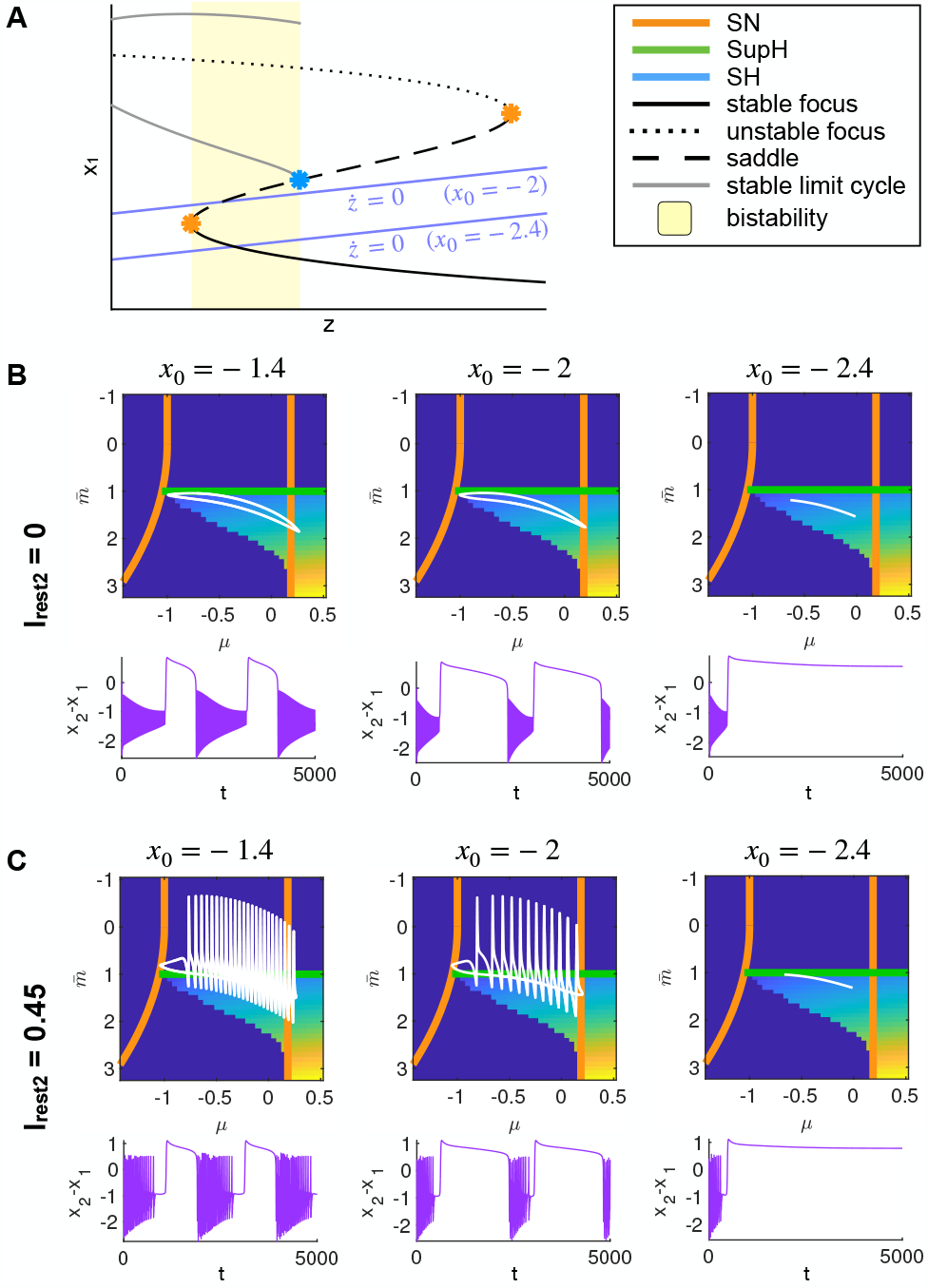
Role of *x*_0_ on the path. **A**. A sketch of how, in hysteresis-loop bursting, *x*_0_ changes the position of the z-nullcline with regards to the resting state branch. **B**. When the intermediate subsystem is not allowed to oscillate (*I*_*rest*2_ *>* 0), different values of *x*_0_ do not change the path but only affect the velocities at which the slow variable evolves when the fast ones are in the interictal or ictal states. **C**. Allowing for intermediate oscillations to modulate the path (*I*_*rest*2_ *>* 0), changing *x*_0_ modifies the amount of spikes in the path (more spikes when the slow variable evolves slower while in the ictal state).

## 4 Discussion

In this work we used a minimal model for SN/SH bursting, that is the DTB bursting model, to investigate a more complex phenomenological model for seizure activity that encapsulates this type of bursting, the Epileptor model. Previous Epileptor’s bifurcation analysis have focused on the two parameters *m* and *x*_0_ [El Houssaini et al., 2020]. Our contribution here is to (i) use the map and movement on the map approach to understand the role of fast parameters on the one hand-side and slow and intermediate variables on the other, (ii) maintain all the parameters of the fast subsystem explicit in the bifurcation analysis to gain insights about their role in shaping the map, (iii) analyze previously described Epileptor’s behaviors placing them in a single map together with new behaviors predicted by the generic model, highlighting the structure present in the dynamic repertoire of the Epileptor.

We started with the fast subsystem of the Epileptor and made all the parameters in it explicit. We identified those that slowly change due to coupling with slower variables, 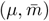, and analyzed their bifurcation diagram, demonstrating the topology to be equivalent to that of the DTB bursting model in the vicinity of the path for SN/SH bursting. Once obtained this map, we illustrated trajectories followed by the full Epileptor model, simulated with choices of parameters from the literature, producing SN/SH and SN/SupH hysteresis-loop bursting and depolarization block [El Houssaini et al., 2015; Jirsa et al., 2014]. In addition to already known classes, we exploited knowledge from the DTB bursting model to change the model’s parameters to produce other two types of hysteresis-loop bursting: SupH/SH and SupH/SupH and to state that no other classes are possible in this map. All the classes identified so far occur in a bistability region in which the resting state fixed point is outside the limit cycle representing the ictal state. This translates in a jump in the baseline of the signal at seizure onset and offset. While this is a very common feature in human seizures, to the point that the presence of a direct current (DC) shift alone has proved indicative of the EZ [Ikeda et al., 1996; Kanazawa et al., 2015], patients exhibits also other types of seizures, including those with no DC shift, that cannot be accounted for, at the moment, by the Epileptor [Saggio et al., 2020].

With regards to the intermediate subsystem, we showed how it modulates the path, causing the fast subsystem to go through a sequence of Hopf bifurcations while moving towards seizure offset and, sometimes, altering the actual offset bifurcation. Finally we commented on the effect on the path of changing *x*_0_, the Epileptor’s parameter used to set the epileptogenicity of brain regions in large-scale Virtual Epileptic Patient models.

Seizures sometimes fail to terminate, evolving into Refractory Status Epilepticus (RSE), a dangerous condition difficult to treat. Two mechanisms for RSE have been proposed in the models we are dealing with and they both involve region V of the map, in which a limit cycle is the only stable attractor. This could help explain why this status is so difficult to reverse. The first mechanism (see El Houssaini et al. [2015] Fig 2) relies on the fact that, in a certain part of region V, the average *x*_1_ activity falls below the z-nullcline so that the behavior of *z* inverts. Now, even if the fast subsystem is not at rest the slow variable promotes movement away from the offset bifurcation. This mechanisms requires an ultra-slow drift to bring the system far enough in region V to enter this regime. The second mechanism (see Saggio et al. [2020] Fig 5 D-I and Fig 14 in Appendix 1) leverages on the presence of a high level of noise in the system, which can occasionally override the slow-variable mechanism and prevent the fast subsystem to reach the offset bifurcation. It could repeatedly approach and fail to reach this bifurcation (in region III) or dwell in region V if an ultra-slow drift is present. Interestingly, only some of the ‘noisy’ SN/SH simulations evolved into RSE, with the rate increasing for higher levels of noise. A similar mechanisms for RSE has been hypothesized in a biophysically-inspired neural mass model [Kramer et al., 2012] and, given the topological equivalence, could be reproduced also with the Epileptor.

The hysteresis-loop bursting mechanism produces periodic autonomous seizures as those observed, for example, in *in vitro* hippocampal preparations. The statistics of the distributions of seizure onset and offset in patients, however, while justifying the use of a feedback mechanism for termination, gives heterogeneous results for seizure iniziation [Suffczynski et al., 2006]. Seizure onset seems to be modulated by a series of mechanisms, spanning several time-scales, such as hormonal, genetic, environmental, sleep-wake cycle and behavior factors [Baud et al., 2018] that may act independently from the fast subsystem, occasionally bringing the latter in regions of the map close to seizure onset. We can thus group the wide range of timescales involved in seizure activity into three main groups, with each group pointing to different meachanisms: fast (faster than ictal length), slow (ictal length) and ultra-slow (slower than ictal length). Fast variables relate to the neuroelectric processes responsible for the generation of oscillations during the seizure (fast oscillations, spike and wave complexes, high frequency oscillations…). Both fast and intermediate Epileptor’s variables fall into this category and have been hypothesized to reflect the activity of glutamatergic and GABAergic cells respectively. Slow variables, such as the Epileptor’s *z*, are those responsible for seizure termination through mechanisms well represented by the feedback loop of the Epileptor model. They can be linked to a variety of processes including ionic currents, metabolic processes, alteration in the intracellular or extracellar environments, neuromodulation, but also the effects of the modulatory effect of some long-range connections, to cite a few. Finally ultra-slow variables are those, already mentioned above, responsible for bringing the system close to seizure onset, but also for transitions between seizure types or even for epileptogenisis. These variables are not modeled in the Epileptor. Both slow and ultra-slow processes are likely related to neurochemical actions [Saggio et al., 2020].

In this work we identified the two fast parameters that are slowly changed by *z* and used part of the unfolding of the DTB singularity to guide the investigation of their bifurcation diagram. If other fast parameters of the Epileptor could be allowed to slowly change, it is possible that a similar map could be obtained for different parameters combinations, which poses a problem at the current state in trying to link the parameters of the two models. For example, in the DTB unfolding, we can obtain a topologically equivalent map also for layers with a fixed *ν* (small and positive). One possibility to start a more robust mapping between the models would be identifying a DTB singularity in the Epileptor. Interestingly, this singularity seems to play a crucial role in the organization of bifurcation diagrams of several neural and neural mass models. Kirst et al. [2015] have demonstrated that conductance based models for neurons contain such a singularity, while Touboul et al. [2011] identified the DTB point in two physiologically inspired neural mass models that have been used in the context of seizures modeling: the Jansen-Rit model and the Wendling-Chauvel. The identification of such singularity, thus, could prove to be a useful tool in analyzing a variety of models and possibly link physiological and phenomenological variables [Depannemaecker et al., 2022]. Another advantage of such an approach is that it would help identify parameters of the Epileptor model that, if allowed to slowly change, could produce other forms of bursting present in the DTB model, for example those without a DC shift that are absent from the current Epileptor’s map.

## 5 Methods

### 5.2 Model

The DTB bursting model presented in Equation 2 is a simplification of the original model from Saggio et al. [2017] that only holds in this portion of the unfolding, in which the two versions of the model behave similarly with regards to bursting. The differences are three. (i) The original model used a 2D map given by the surface of a sphere of small radius centered around the DTB singularity, while here, with the only goal of simplifying the description of the model, we used a portion of the unfolding on a layer obtained with a small and fixed *μ*_2_. (ii) As a consequence of the previous choice, we could use simple segments as paths for bursting, whereas in the original model the paths were arcs of great circles. (iii) For the slow dynamics, the original model uses a more complex description that ensures that the model works even if the resting state is inside the limit cycle, which never occurs in this portion of the map.

As it is used only with an explanatory intent, Fig 1B is produced with the original model.

### 5.2 Bifurcation analysis

Bifurcation curves in Fig 1B are obtained using Matcont [Dhooge et al., 2003]. The bifurcation analysis of the Epileptor has been done analytically and using the symbolic Matlab toolbox with regards to local bifurcation.

For the SH bifurcation we performed simulations of the fast subsystem (using Matlab function ‘ode45’) for the different combinations of parameters values as shown in Fig 4, using the fixed point that is not the resting state plus *ϵ* = 0.01 as initial conditions. For the Epileptor, we simulated 600 s, removed the first 300 s to avoid the transient behavior and used the remaining to compute the amplitude and frequency of *x*_1_. For the amplitude we took the difference between the maximum and minimum of the timeseries; for the frequency we used Hann window and then applied Discrete Fourier Transform. We computed amplitude and frequency of the limit cycle for the DTB model with the same procedure (as in [Saggio et al., 2017]), simulating 2000 s and removing the first 500 s.

#### 5.2.1 Simulations

All simulations are performed without noise, using Matlab function ‘ode15s’ with maximum integration step 0.01. Parameters of the model are set as in Jirsa et al. [2014] unless otherwise stated, except for *x*_0_ = −2 for improved visualization purposes. As described in Fig 7, this does not alter the onset/offset patterns that are the main focus of this work.

